# Emergence of B2-ST131-C2 and A-ST617 *Escherichia coli* clones producing both CTX-M-15- and CTX-M-27 and ST147 NDM-1 positive *Klebsiella pneumoniae* in the Tunisian community

**DOI:** 10.1101/713461

**Authors:** Amel Mhaya, Rahma Trabelsi, Dominique Begu, Sabine Aillerie, Fatima M’zali, Slim Tounsi, Radhouane Gdoura, Corinne Arpin

## Abstract

CTX-M ESBLs have been increasingly reported from human enterobacteria isolates in Tunisia. NDM-1 carbapenemase was recently reported in isolates from Tunisian Hospitals. During a 2-month period (December 2017 to January 2018), we have collected 23 ESBL-producing *Enterobacteriaceae* (ESBL-E) from urines of patients hospitalized in three health care facilities and from community, situated in two distinct Tunisian geographical regions. They were divided into 15 *Escherichia coli* and eight *Klebsiella pneumoniae*. The aim of this study was to characterize ESBL-E with regard to their β-lactamase content and their epidemiological relationship. The results indicated a high rate (47%) of *E. coli* producing both CTX-M group-1 and −9. For the first time, we demonstrated the presence of *E. coli* having concomitantly CTX-M-15 and CTX-M-27 which belong to two sequences types (ST), *ie.* A-ST617 (2 isolates) and B2-ST131 subclade C2 (2 isolates). All four *E. coli* isolates carried a multireplicon IncF with an identical allelic combination, F31:A4:B1. This study reports also the first description of *K. pneumoniae* belonging to the clone ST147 carrying the carbapenemase NDM-1 in the Tunisian community. In conclusion, our data confirms the need for monitoring the resistance to extended-spectrum cephalosporins and to carbapenems among enterobacteria in Tunisia.

## INTRODUCTION

Resistance to extended-spectrum cephalosporins (ESC) is widespread among *Enterobacteriaceae* species and is mainly due to the production of extended-spectrum ß-lactamases (ESBLs). The dissemination of ESBLs is become a growing concern since they are considered as a major cause of morbidity and mortality. Up to the end of the 1990s, TEM- and SHV-type ESBLs were mainly produced by the *Klebsiella pneumoniae* and *Enterobacter* spp. responsible for nosocomial infections (1, 2). Since 2000s, CTX-M ESBLs have gained prominence mainly in *Escherichia coli* and *K. pneumoniae* strains and are now considered as pandemic enzymes. Many *bla*_CTX-M_ variants exist, but they can be divided into two main clusters (groups −1 and −9); each cluster of CTX-M genotypes has a corresponding progenitor gene sharing homology with different environmental *Kluyvera* spp. from which *bla*_CTX-M_ genes originated. Among CTX-M enzymes, CTX-M-15 belonging to group 1, has currently been the most frequent all over the world (1, 3)

The majority of CTX-M type ESBL-associated *E. coli* infections is often concentrated within specific extraintestinal pathogenic *E. coli* (ExPEC) lineages belonging to highly virulent phylogenetic group B2, and recognizable by their sequence type (ST). The *bla*_CTX-M-15_ is the dominant ESBL gene in the virulent *E. coli* ST131 clone, in particular in the subclade ST131-C2 (4). However other genetically divergent CTX-M genes also occur in this ST, such as *bla*_CTX-M-14/14-like_ variants in Canada, China, and Spain (5, 6). A subclade of *E. coli* ST131 producing the CTX-M-27 (group 9 enzyme, CTX-M-14 variant), named ST131-C1-M27 has emerged more recently (7).(8)

The *bla*_CTX-M_ genes have mainly been disseminated via multiple horizontal gene transfer events (9), and they are often located on conjugative plasmids, especially multireplicon IncFII plasmids additionally harboring FIA/FIB replicons, which also carry other resistance genes (10). The important dissemination of ESBLs has compromised the use of ESC and promoted the use of carbapenems and consequently favored the diffusion of carbapenemase-producing isolates such as the metallo-enzyme NDM-1 (11). In Tunisia, several studies reported the presence of ESBLs in *Enterobacteriaceae* (12, 13). The presence of *K. pneumonia-*NDM-1 positive strains were found in Tunisian Hospitals and suspected to be also present in community (30, 31). Thus, the need for systematic monitoring of ESBL- and carbapenemase-mediated resistance in Hospital and community settings is important.

In the present work, we studied the characteristics of ESBL-producing enterobacteria collected from urines of patients in Tunisia. During a 2-month period, we collected 200 urine samples that gave positive bacterial cultures, and identified 170 enterobacteria. Among them, strains were randomly collected from patients of a Urology unit of a polyclinic situated in Sfax, and compared to isolates obtained from two distant Tunisian health care facilities. ESBL-producing enterobacteria (ESBL-E) collected in the Sfax community were also included. The strains were characterized with regard to antibiotic resistance, β-lactamase content, plasmid-mediated ESBL, and clonality (pulsotype profiles and MultiLocus Sequence Typing).

## RESULTS

### Prevalence of ESBL-producing enterobacteria in Tunisian health care facilities

A collection of urine positive culture samples selected as a random sampling was recovered from December 2017 and January 2018. They came from three health care facilities situated in two Tunisian geographic regions, Sfax and Tunis; both locations are ca. 270-km away. One of them (polyclinic, H1) is situated at Sfax with samples mainly collected from patients of an Urology unit. The other health care facilities were located in Tunis (H2, regional Hospital and H3, psychiatric clinic). Thus, a total of 118 enterobacteria were recovered which corresponded to non-repetitive isolates (collected from different patients) (Table 1). A total of eight ESBL-E were identified as confirmed by phenotypic and molecular characterizations, leading to ESBL-E prevalence of 6.25%, 11.1%, and 3.7% in H1, H2 and H3, respectively (Table 1). Regarding the rate of ESBL-producing *E. coli*, the data were as following: 7.5% (H1), 20% (H2) and 6.6% (H3). ESBL-E were collected from patients with a sex ratio of 1.

**Table 1.** Distribution of ESBL-producing enterobacteria among a collection of 170 enterobacteria positive urine samples

| Strain | Heath care facility<sup>(a,b)</sup> |  |  | Community<sup>(a, c)</sup> |  |
| --- | --- | --- | --- | --- | --- |
|  | H1 | H2 | H3 | CA | CB |
| *E. coli* | 3 (n= 40) | 3 (n=15) | 1 (n=15) | 1 (n=10) | 7 (n=30) |
| *K. pneumoniae* | 1 (n=20) | 0 (n=10) | 0 (n=10) | 3 (n=5) | 4 (n=5) |
| *E. cloacae* complex | 0 (n=4) | 0 (n=2) | 0 (n=2) | 0 (n=0) | 0 (n=2) |

<sup>(a)</sup>, Number of ESBL-E (total strains number is indicated in parenthesis)
<sup>(b)</sup> Strains provided from Health care facilities (H1 to H3), <sup>(c)</sup> and from Community settings (CA and CB Laboratories).

### Characterization of ESBL-producing enterobacteria

During the same period, 52 multidrug resistant enterobacteria were collected from urines of community subjects at Sfax. The aim was to determine the ESBL profile and epidemiological relationship of enterobacteria in the Tunisian community setting and to compare with those isolated from health care facilities. From two private Laboratories (CA and CB), 15 ESBL-E including seven ESBL-producing *K. pneumoniae*. Thus, a total of 23 ESBL-E were obtained in our survey (8 from health care facilities and 15 from community) divided into 15 *E. coli* and eight *K. pneumoniae* (Table 1).

As indicated in Table 2, all 15 ESBL-producing *E. coli* gave positive PCR amplifications with specific primers of the *bla*_CTX-M_ gene. Six and four isolates respectively, also exhibited OXA-1 like and TEM-1-like ß-lactamases, including one which carried both enzymes. Surprisingly among the 15 CTX-M-producing *E. coli*, almost the moiety (7 strains, 47%) gave PCR positive amplifications for both groups of CTX-M (group 1 and group 9), five *E. coli* carried a single CTX-M group 1 enzyme and the three remaining, a CTX-M group 9.

**Table 2.**
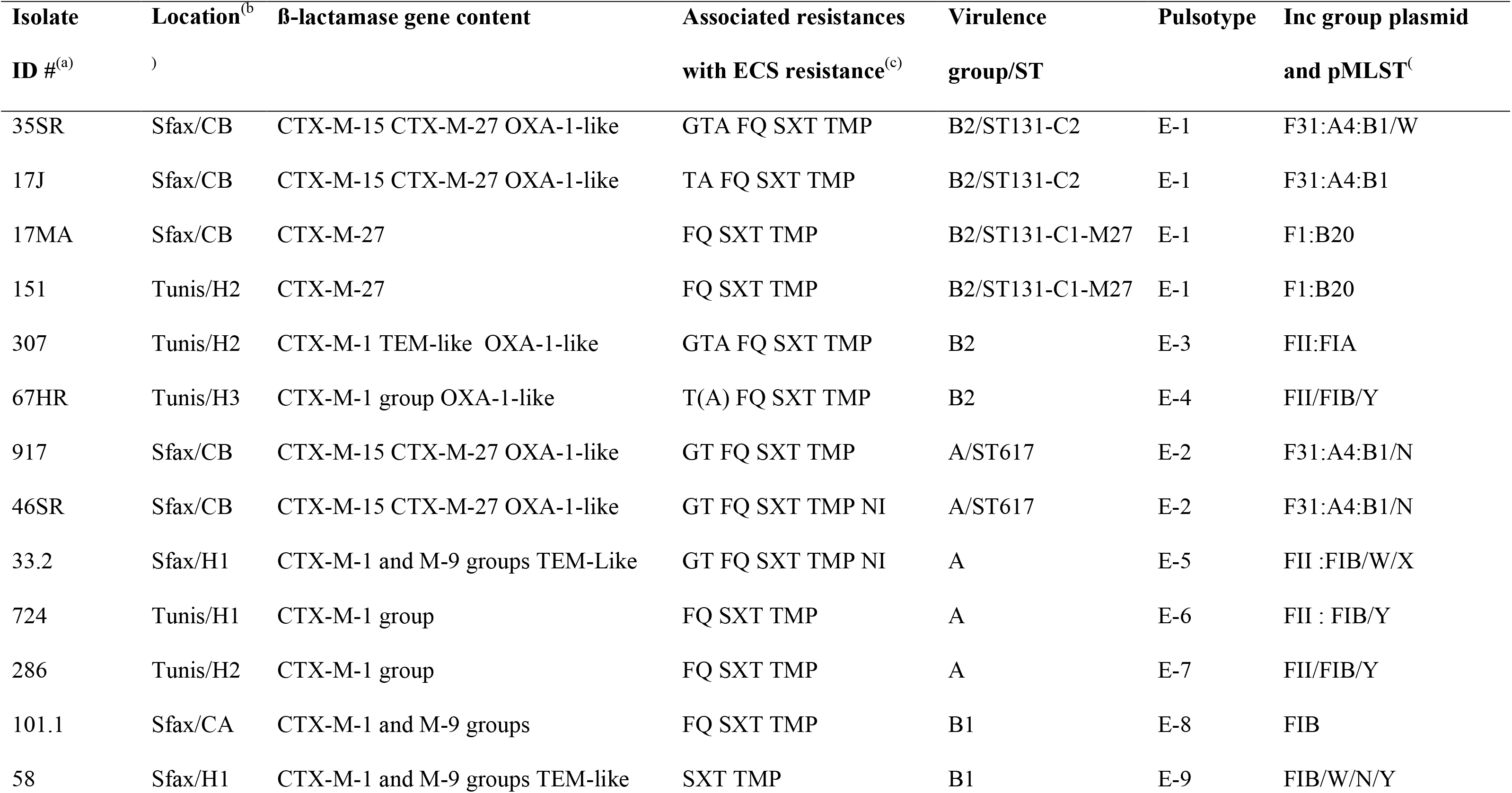
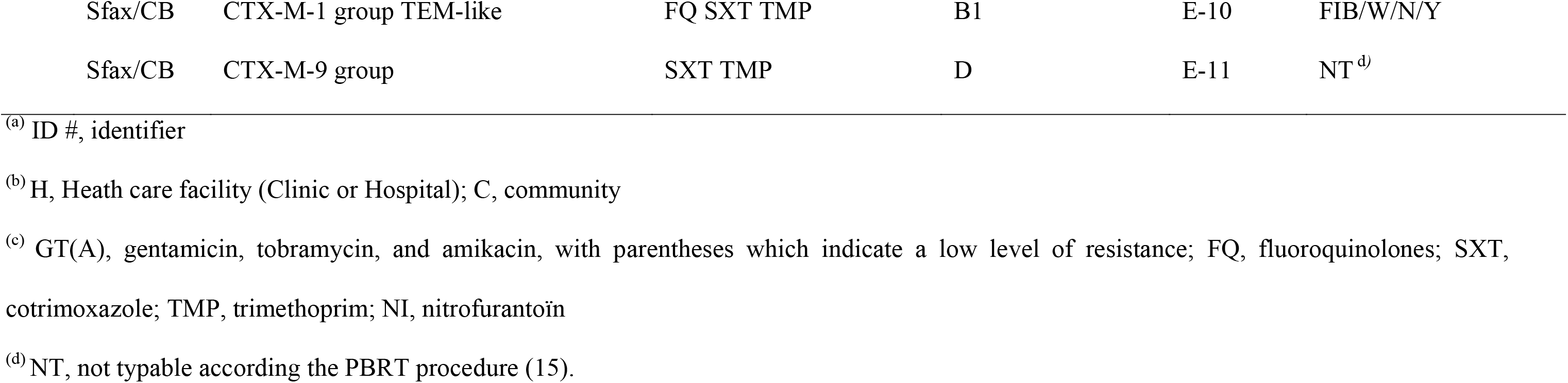
Characteristics of the 15 ESBL-producing *E. coli* isolated from our survey

| Isolate<br>ID #<sup>(a)</sup> | Location<sup>(b)</sup><br>) | β-lactamase gene content | Associated resistances<br>with ECS resistance<sup>(c)</sup> | Virulence<br>group/ST | Pulsotype | Inc group plasmid<br>and pMLST<sup>(c)</sup> |
| --- | --- | --- | --- | --- | --- | --- |
| 35SR | Sfax/CB | CTX-M-15 CTX-M-27 OXA-1-like | GTA FQ SXT TMP | B2/ST131-C2 | E-1 | F31:A4:B1/W |
| 17J | Sfax/CB | CTX-M-15 CTX-M-27 OXA-1-like | TA FQ SXT TMP | B2/ST131-C2 | E-1 | F31:A4:B1 |
| 17MA | Sfax/CB | CTX-M-27 | FQ SXT TMP | B2/ST131-C1-M27 | E-1 | F1:B20 |
| 151 | Tunis/H2 | CTX-M-27 | FQ SXT TMP | B2/ST131-C1-M27 | E-1 | F1:B20 |
| 307 | Tunis/H2 | CTX-M-1 TEM-like OXA-1-like | GTA FQ SXT TMP | B2 | E-3 | FII:FIA |
| 67HR | Tunis/H3 | CTX-M-1 group OXA-1-like | T(A) FQ SXT TMP | B2 | E-4 | FII/FIB/Y |
| 917 | Sfax/CB | CTX-M-15 CTX-M-27 OXA-1-like | GT FQ SXT TMP | A/ST617 | E-2 | F31:A4:B1/N |
| 46SR | Sfax/CB | CTX-M-15 CTX-M-27 OXA-1-like | GT FQ SXT TMP NI | A/ST617 | E-2 | F31:A4:B1/N |
| 33.2 | Sfax/H1 | CTX-M-1 and M-9 groups TEM-Like | GT FQ SXT TMP NI | A | E-5 | FII :FIB/W/X |
| 724 | Tunis/H1 | CTX-M-1 group | FQ SXT TMP | A | E-6 | FII : FIB/Y |
| 286 | Tunis/H2 | CTX-M-1 group | FQ SXT TMP | A | E-7 | FII/FIB/Y |
| 101.1 | Sfax/CA | CTX-M-1 and M-9 groups | FQ SXT TMP | B1 | E-8 | FIB |
| 58 | Sfax/H1 | CTX-M-1 and M-9 groups TEM-like | SXT TMP | B1 | E-9 | FIB/W/N/Y |
| 17G | Sfax/CB | CTX-M-1 group TEM-like | FQ SXT TMP | B1 | E-10 | FIB/W/N/Y |
| 18AF | Sfax/CB | CTX-M-9 group | SXT TMP | D | E-11 | NT<sup>d)</sup> |

<sup>(a)</sup> ID #, identifier
<sup>(b)</sup> H, Health care facility (Clinic or Hospital); C, community
<sup>(c)</sup> GT(A), gentamicin, tobramycin, and amikacin, with parentheses which indicate a low level of resistance; FQ, fluoroquinolones; SXT, cotrimoxazole; TMP, trimethoprim; NI, nitrofurantoin
<sup>(d)</sup> NT, not typable according the PBRT procedure (15).

Among the eight ESBL-producing *K. pneumoniae*, all strains were collected from community subjects at Sfax, except one strain (ID, # 115) which provided from H1. Three and four strains were collected from Laboratories CA and CB, respectively. All the strains contained a group 1 CTX-M. Six of them also harbored the *bla*_OXA-1-like_ gene, including four together with *bla*_TEM-1-like_. A strain provided from the Laboratory CB showed a carbapenem resistance phenotype and harbored the metallo-enzyme, NDM-1 as confirmed by sequencing. This isolate (ID, # 18TA) had been firstly classified as *Enterobacter spp.* by biochemical tests (*i.e.* API 10S gallery including a negative-urease test) gave *K. pneumoniae* identification by MALDI-TOF. PCR amplification and sequencing of rDNA 16S confirmed the *K. pneumoniae* identification. The CTX-M group 1 present in this isolate (ID, #18TA) was furthermore characterized and showed the presence of the CTX-M-15 ß-lactamase.

### Strain epidemiological typing analysis

The 15 CTX-M-producing *E. coli* mainly belonged to phylogenetic group B2 (40%), followed by the groups A (33.3%), B1 (20%) and D (6.7%) (Table 2). Considering a cluster defined at a 68% similarity level (13), the 15 *E. coli* were assigned to 11 distinct PFGE patterns (pulsotypes E-1 to E-11, Figure 1 and Table 2). Four isolates of subgroup B2 divided into a similar pulsotype E-1 (ID# 35SR; 17J, 17MA and 151, Figure 1, lanes 1 to 4) and two isolates of subgroup A (ID# 917 and 46SR) exhibited an identical profile E-2. To point out the phylogenetic relationship of the six strains, MLST typing were performed and showed that the two isolates of E-2 profile can be including in the sequence type A-ST617 and the four E-1 profile belonged to the clone B2-ST131 (Table 2). Among the four later isolates, two carried a CTX-M-27, and the two other one exhibited both CTX-M-15 and CTX-M-27. In literature, the ST131-B2 *E. coli* having CTX-M-27 enzyme are often situated in the subclade C1-M27 (7). To verify the lineage of the ST131 *E. coli* carried both CTX-M-15 and/or CTX-M-27, we have included in our study two strains of B2-ST131 collected in a previous Tunisian survey and performed from December 2014 to February 2015 (14): one with CTX-M-15 (ID# 59) and the other one with CTX-M-27 (ID# 89). The isolates with the single ESBL, CTX-M-27 (ID# 17MA and 151 and the control strain ID# 89) belonged to the clade C1-M27. The two strains which exhibited both CTX-M-15 (group 1) and CTX-M-27 (group 9) belonged to the subclade ST131-C2, as that found for the control isolate (ID# 59) which only harbored CTX-M-15. The *K. pneumoniae* 18TA isolate carrying both ESBL (CTX-M-15) and carbapenemase (NDM-1) belonged to ST147.

**Figure 1.**
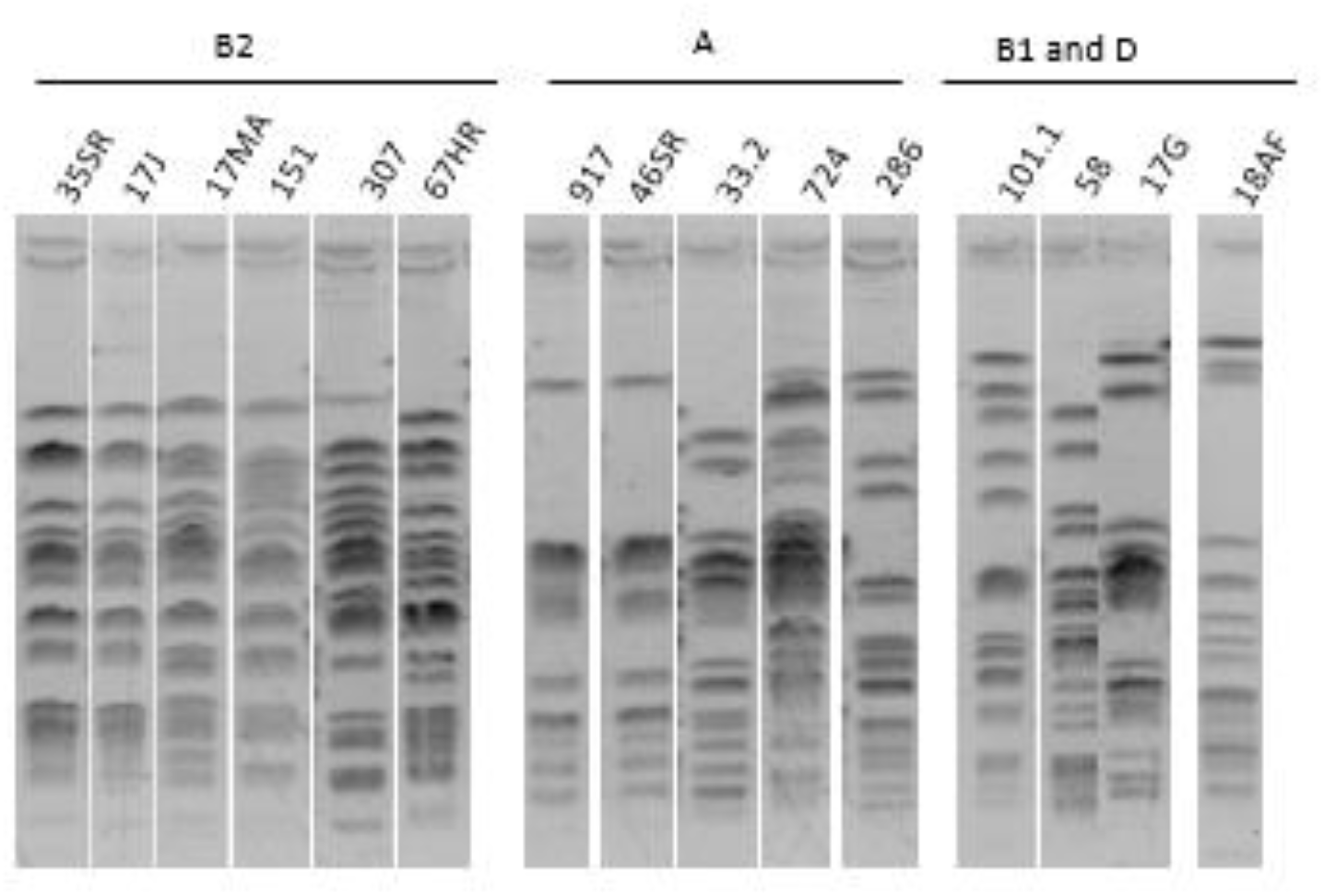
Pulsotypes of 15 CTX-M-producing *E. coli.* The pulsotypes were classified according their phylogroups B2, A, B1 and D, and their pulsotypes.: E-1 (lanes 1 to 4); E-3 and E-4 (lane 5 and 6, respectively); E-2 (lanes 7 and 8); E-5 to E-11 (lanes 9 to 15, respectively).

### Antibiotic resistance patterns and plasmid typing

All ESBL-producing *E. coli* strains showed decreased susceptibility to ß-lactams, except to cephamycins (*ie.* cefoxitin) and carbapenems. All remained susceptible to fosfomycin, and two isolates were resistant to nitrofurantoin (Table 2). All *E. coli* were resistant to co-trimoxazole (sulfamethoxazole-trimethoprim) and all, except two isolates of phylogenetic groups B1 and D, were resistant to fluoroquinolones (Table 2). They exhibited different aminoglycoside phenotypes (resistance to gentamicin, tobramycin and amikacin, 26%, 47% and 20%, respectively) (Table 2). Among the *K. pneumoniae* strains, they exhibited various profiles to tested antibiotics (Table 3). A single isolate (ID# 18TA) with the NDM-1 enzyme showed a diminution of diameter to imipenem which was restored by addition of EDTA. Plasmids of *E. coli* were characterized by PCR-based replicon typing (15). Most of them were assigned to a multi-replicon replicon IncFII (Table 2). A sub-typing scheme of this replicon using a FAB formula (FII:FIA:FIB) was performed for the six *E. coli* isolates of pulsotypes E-1 and E-2. Thus, four *E. coli* including the two isolates of pulsotype E-2 and two other ones (E-1) ascribed to lineages A-ST617 and B2-ST131-C2, respectively harbored the formula replicon: F31:A4:B1 (Table 2). Other plasmids were also found in these epidemic isolates, *ie.* IncN, in the isolates ID# 917 and ID# 46SR and IncW, in the isolate ID# 35SR. None of ESBL genes in the four *E. coli* was transferred by mating out assays. The two other *E. coli* B2-ST131-C1-M27 are contained on an IncFII plasmid having the formula F1:B20. In contrast to six tested *E. coli*, all *bla* genes and associated co-resistances present in *K. pneumoniae* 18TA were transferred by conjugation in Az^R^ *E. coli* C600.

**Table 3.** Characteristics of the eight ESBL-producing *K. pneumoniae*

| Isolate ID #<sup>(a)</sup> | Sfax<br>location<sup>(a)</sup> | β-lactamase gene content<sup>(b)</sup> | Associated resistances<br>with ESC resistance<sup>(a)</sup> |
| --- | --- | --- | --- |
| 115 | H1 | CTX-M-1 group OXA-1-like | FQ |
| 96.1 | CA | CTX-M-1 group OXA-1-like | SXT |
| 57 | CA | CTX-M-1 group OXA-1-like TEM-like | G FQ SXT |
| 39/GB | CA | CTX-M-1 group OXA-1-like TEM-like | - |
| 18TA (ST147)<sup>(c)</sup> | CB | CTX-M-15 NDM-1 OXA-1-like TEM-like | GTA FQ SXT TMP NI |
| 48SR | CB | CTX-M-1 group OXA-1-like | GTA FQ SXT TMP NI |
| 17MZ | CB | CTX-M-1 group TEM-like | G FQ |
| 18CT | CB | CTX-M-1 group OXA-1-like TEM-like | FQ |

<sup>(a)</sup> Abbreviations as defined in Table 2
<sup>(b)</sup> All the *K. pneumoniae* strains gave a positive PCR with SHV-specific primers which probably corresponds to the amplification of the β-lactamase chromosomal natural-species gene.
<sup>(c)</sup> Sequence type of the isolate 18TA is given

## DISCUSSION

The presence of CTX-M ESBLs and of carbapenemases such as NDM-1 from human enterobacteria isolates has been increasingly reported worldwide, including Tunisia.(1, 11, 16). In this 2-month survey, the incidence rate of ESBL-producing *Enterobacteriaceae* from urine collected from patients hospitalized in three health care facilities from two geographical areas of Tunisia, ranged from 11.1% to 3.7%. Data are consistent with those described in literature which showed variable incidence rates depending of risk factors of ESBL-E acquisition (17, 18). Few studies including molecular analysis ESBL-E from community-acquired infections were reported in Tunisia (16). However, several reports from outpatients and from enterobacteria fecal carriage performed at Hospital admission have also underlined the presence of CTX-M-producing enterobacteria in the Tunisian community setting (14).

Nevertheless, the main finding of our work is the high rate of different lineages of extraintestinal pathogenic *E. coli* which produced concomitantly CTX-M ESBLs of group 1 and group 9. Some studies mainly on the Asian continent mentioned *E. coli* or *K. pneumoniae* isolates with both CTX-M-55 and CTX-M-14 or CTX-M-15 and CTX-M-14 (19, 20). To the best of our knowledge, this study is the first description of *E. coli* with both CTX-M-15 and CTX-M-27. The two enzymes, with a substitution Asp240 to Gly in their active site are known to increase their hydrolytic activity toward ceftazidime, with respect to CTX-M variants without this substitution (21, 22); suggesting than a high ceftazidime selection pressure could favor the emergence of such isolates. Thus, we report the presence of the two enzymes in at least four fluoroquinolone-resistant *E. coli* isolates belonging to two different STs. Two of them were associated to the pandemic and virulent clone ST131, belonging to the phylogenetic group B2. Recent studies using whole-genome sequencing analysis revealed that ST131 consists of different lineages or clades: A/H41, B/H22, and C/H30; with the H number which refers to the *fimH* allele which mostly corresponds to each clade (4, 23, 24). Since the 2000s, clade C has become the most dominant lineage among ST131 isolates (7, 9). The clade C mainly consists of the fluoroquinolone resistant subclades C1 and C2 isolates, with subclade C2 (also known as H30Rx) which is often associated with *bla*_CTX-M-15_ gene (4). Recently, a C1 subclade associated with the *bla*_CTX-M-27_ gene, named C1-M27, has been described on the Asian continent and also reported with an alarming increase in Europe.(6, 7, 25). The presence of 025b-H30x ST131 *E. coli* having CTX-M-27 was described in Tunisia from human samples (14), and also from animal origin and wastewater (26), suggesting a such emergence can also happen in this country. Using the recently C subclades identification by PCR amplifications, we confirmed that the CTX-M-27-producing *E. coli* ST131 belongs to the subclade C1-M-27, similarly to that (isolate ID # 89) collected from a Tunis Hospital during a survey in 2014-2015 (14). By using this PCR procedure, we also showed that two *E. coli* ST131 provided from Sfax community which carried both CTX-M-15 and CTX-M-27. These strains were phylogenetically closer to subclade C2 than subclade C1, similarly to the control *E. coli* isolate with the single CTX-M-15 (ID #, 59) (14). Thus, our data suggest an additional acquisition of CTX-M-27 in the pandemic subclade C2. Both CTX-M-15 and M-27 enzymes have also been found in another clonal strain of *E. coli* which belonged to the phylogenetic group A and ST617 (clonal complex 10, CC10). CTX-M-15-producing *E. coli* strains ST617 were also widespread internationally and is described in relation to human infections as well as in commensal *E. coli* strains isolated in humans, animals and food samples (27).

The spread of *bla*_CTX-M_ genes is largely mediated by conjugative plasmids, and especially IncF one which often carry *bla*_CTX-M-15_ gene or the increasingly *bla*_CTX-M-27_ in *E. coli* ST131; these plasmids likely contributing to the success of this ST (28, 29). In our study, the *E. coli* strains A-ST617 and subclade C2 B2-ST131 exhibited the same IncF multireplicon combination (FII:FIA:FIB), *ie.* F31:A4:B20, suggesting a possible plasmid transfer. However, in our laboratory experiments, no conjugation was observed. A same IncF combination (*ie*. F31:A4:B1) plasmid carrying *bla*_CTX-M-15_ has regularly been identified in ST617 and ST131 isolates in Tunisia and different countries worldwide and from various samples (30). It will be interesting to perform the whole-sequencing of our plasmids to analyze their genetic relationship and evolution and to know if the acquisition of CTX-M-27 in a CTX-M-15-mediated plasmid in ST617 and S131 could be the result of independent events. These data could contribute to enhance the knowledge about the ST131 evolution and their associated replicons. A second F-type plasmid (F1:B20) was identified among two other strains ST131 belonging to the clade C1-M27. In literature, the F1:A2:B20 plasmid is strongly associated with the C1-M-27 clade isolates (29); the F1:B20 without FIA replicon were reported in Spain (https://pubmlst.org/plasmid/primers/incF.shtml).

Regarding the important number of CTX-M-producing *K. pneumoniae* collected from Tunisian community, our results corroborate previous studies where ESBL positive *K. pneumoniae* are often isolated from this setting (14). NDM-positive *K. pneumoniae* ST147 lineage strains are also relatively common in multiple countries across several continents, almost all of which were isolated from humans (11). Recently, reports indicated the presence of ST147 NDM-1-producing *K. pneumoniae* in Tunisian Hospitals, but no data indicate its spread in community (8, 31). We report the presence of this clone in the community context; its prevalence may be underestimated.

In conclusion we report, for the first time, not only the presence of CTX-M-15 and M-27 in two different STs of *E. coli*, including ST131 which is a predominant lineage among ExPEC and which plays a major role in the worldwide dissemination of CTX-M-producing *E. coli*, but also the presence of the ST147 NDM-1-producing *K. pneumoniae* clone in Tunisian community. The monitoring of these potential emergent clones is urgently needed, and preventive programs to limit their dissemination should be implemented.

## MATERIAL AND METHODS

### Strains collect and identification of enterobacteria

Between December 2017 and January 2018, a total of 200 urine positive cultures were collected. They provided from patients of three heath care facilities, including one (H1, of about 60-bed) situated at Sfax and two other ones located at Tunis. Urine samples have also been recovered from community patients from two private biological Laboratories at Sfax (Laboratories CA and CB). Enterobacteria identification was done by conventional methods including biochemical tests (BioMerieux API 10S) and confirmed by Matrix-Assisted Laser Desorption Ionization-Time-Of-Flight/Mass Spectrometry (MALDI-TOF/MS, Brüker Daltonics). One enterobacteria gave a discordance result between biochemical tests and MALDI-TOF, was furthermore analyzed by rDNA16S PCR and sequencing.

### Antibiotic susceptibility testing

The antibiotic susceptibility patterns of enterobacteria (and the transconjugant of *K. pneumoniae* isolate 18TAs) were determined by the disk diffusion method in Mueller-Hinton (MH, Bio-Rad, Marnes-la-Coquette, France) agar by using the antibiotics as following: ampicillin, amoxicillin-clavulanic acid, ticarcillin, ticarcillin-clavulanic acid, piperacillin, piperacillin-tazobactam, mecillinam, temocillin, cefalexine, cefoxitin, cefixime, cefuroxime, ceftriaxone, ceftazidime, cefepime, aztreonam, imipenem, meropenem, ertapenem, gentamicin, tobramycin, amikacin, acid nalixidic, ciprofloxacin, ofloxacin, colistin, co-trimoxazole, trimethoprim, fosfomycin, nitrofurantoin (Bio-Rad). Results were interpreted according criteria to the European Committee on Antimicrobial Susceptibility Testing (https://www.sfm-microbiologie.org/2019/01/07/casfm-eucast-2019/). Isolates were screened for ESBL production by double-disk synergy test (2). Isolates showing increased resistance to carbapenems, a susceptibility of EDTA test was conducted for metallo-carbapenemase detection.

### Molecular characterization of ß-lactamase genes

After extraction, total DNA of all ESC-resistant enterobacteria was screened for the search of ß-lactamases which included multiplex PCR by amplifications for types TEM-, SHV- and OXA-1-like enzymes and a multiplex PCR for groups −1, −2, −9, −18, −25 CTX-M ß-lactamases (32). In addition, plasmid-mediated AmpC β-lactamase (*i.e.* the ACC, FOX, MOX, DHA, CIT) and carbapenemases (OXA-48 like, KPC, GES, VIM, IMP, NDM) genes were also searched by using a procedure described elsewhere (33, 34). According results, entire genes for selected strains were amplified and sequenced to identify *bla*_CTX-M_ and *bla*_NDM_ subtypes using specific PCR primers known from their close genetic environment (35, 36). The custom-made DNA sequences (Eurofins) were analyzed using the Basic Local Alignment Search Tool (BLAST, http://www.ncbi.nlm.nih.gov/BLAST/).

### Molecular typing of enterobacteria strains

The clonal relationship among *E. coli* and *K. pneumoniae* strains was first investigated after *Xba*I enzyme digestion and pulsed-field gel electrophoresis (PFGE) analysis using the CHEF DRIII apparatus (Bio-Rad) as previously described (35). *E. coli* isolates were assigned to the phylogenetic groups A, B1, B2, or D using a PCR strategy with specific primers for *chuA*, *yjaA*, and *TspE4*.C2 determinants as reported elsewhere (37). Multilocus Sequence Typing (MLST) was carried out for *E. coli* strains with similar pulsotypes and for the NDM-1-producing strain of *K. pneumoniae*. All the amplicons were sequenced and compared with the sequences deposited in the MLST database to know the specific allele combination and the sequence type (ST) for *E. coli* (http://enterobase.warwick.ac.uk/species/ecoli/allele_st_search) and and *K. pneumoniae* (http://www.pasteur.fr/mlst/). Furthermore, we used the specific PCR procedure described by Matsumura *et al.* for clades and subclades assignation of *E. coli* ST131 (7).

### Plasmid analysis

Transferability of ESBL-encoding genes was studied by mating-out assays by using an azide-resistant (Az^R^) mutant of *E. coli* C600 as recipient strain. Selection was performed on MH agar plates supplemented with ampicillin (100 mg/L) or cefotaxime (4 mg/L) and sodium azide (300 mg/L). Plasmid replicons were identified by using the PCR-based replicon typing method (PBRT) (15). The IncF group incompatibility plasmids were further characterized using the IncF replicon typing scheme (https://pubmlst.org/plasmid/primers/incF.shtml).

## ACKNOWLEDGMENTS

We thank Nathalie Peyron for her technical contribution and Pr. Véronique Dubois who provided us with the two CTX-M-producing *E. coli* (ID # 59 and 89) collected from a previous study. This work was supported by the CNRS and University of Bordeaux.

A. M has a fellowship from the Ministère de l’Enseignement Supérieur et de la Recherche Scientifique in Tunisia.

We have no conflicts of interest to declare.

